# Optically active, paper-based scaffolds for 3D cardiac tissue engineering

**DOI:** 10.1101/2024.03.01.582954

**Authors:** F. Guo, S. Jooken, A. Ahmad, W. Yu, O. Deschaume, W. Thielemans, C. Bartic

**Affiliations:** Laboratory for Soft Matter and Biophysics, Department of Physics and Astronomy, KU Leuven, 3001 Leuven, Belgium; Sustainable Materials Lab, Department of Chemical Engineering, KU Leuven, campus Kulak Kortrijk, Etienne Sabbelaan 53, Kortrijk 8500, Belgium; Department of life science technologies, Imec, Kapeldreef 75, 3001, Leuven, Belgium

## Abstract

In this work, we report the design and fabrication of a light-addressable, paper-based, nanocomposite scaffold for optical modulation and read-out of in vitro grown cardiac tissue. The scaffold consists of paper cellulose microfibers functionalized with gold nanorods (GNRs) and semiconductor quantum dots (QDs), embedded into a cell-permissive collagen matrix. The GNRs enable cardiomyocyte activity modulation through local temperature gradients induced by near-infrared (NIR) laser illumination, with the local temperature changes reported by the temperature-dependent QD photoluminescence (PL). The micrometer size paper fibers promote the tubular organization of HL-1 cardiac muscle cells, while the NIR plasmonic stimulation modulates reversibly their activity. Given its high spatial resolution, NIR modulation offers an excellent alternative to electrode-based methods for cell activity modulation and is more compatible with 3D tissue constructs. As such, optical platforms based on nanocomposite scaffolds will have a significant impact on the progress of drug screening, toxicity studies, and heart disease modeling.

## Introduction

Paper-based materials attracted large attention in 3D tissue engineering field, due to their various advantages, such as availability, cost-effectiveness, biocompatibility, and their fiber-based porous microstructure.^[1–3]^ Paper-based materials have been used as cell scaffolds to develop in vitro models, including cardiovascular, ^[4–6]^ liver^[7,8]^ and lung^[9]^ disease models, for disease investigation and drug screening. The fiber density, size, and chemical composition were all found to play important roles in supporting cellular proliferation.^[6,10]^ The cellulose fibers in paper can be used for cell culturing as such^[11,12]^, or in combination with a hydrogel^[13]^, owing to the effective wicking of the paper. Human-induced pluripotent stem cell-derived cardiomyocytes seeded on Matrigel coated paper fiber microstructures formed cell sheets, expressing cardiac specific markers and contractile activity for up to 3 months in vitro.^[6]^ This study demonstrated the potential of using paper-based scaffolds for 3D cardiac tissue modeling, but it still needs further improvements to reproduce the cell morphology and achieve maturation of cardiac tissue.

In this work, we nanoengineered a paper-based scaffold made from the Thorlabs lens cleaning tissue paper to support in vitro 3D cardiac culture. The nanoparticle-functionalized microfiber network in the paper guides HL-1 cardiac muscle cells to organize into functional tubular constructs in 3D and allows synchronizing their electrophysiological activity. The local modulation of the cardiac electrophysiological activity was achieved through gold nanorods (GNRs) mediated plasmonic heating under NIR illumination. At the same time, the local temperature is monitored by the quantum dot (QD) fluorescence readout. This is the first time that NIR-induced thermoplasmonic heating of GNRs for the modulation of cell beating rate is reported in a 3D fibrous scaffold.

Cells sense temperature gradients and adjust their functions (e.g., metabolism, growth, electrophysiological activity) accordingly. Externally applied thermal stimuli have been used to tune cell adhesion,^[14]^ migration,^[15]^ differentiation^[16]^ and electrical activity.^[17,18]^ Plasmonic heating of membrane-targeted GNPs has for instance been shown to evoke significant increases in electrical activity in rat primary auditory neurons,^[19]^ trigger membrane depolarization of dorsal root ganglion neurons and mouse hippocampal neurons, resulting in action potential (AP) firing,^[20]^ as well as increase neurite lengths and the possibility to elicit intracellular calcium transients in NG108-15 neuronal cells.^[21,22]^

In cardiac tissue engineering, the photothermal effect has been investigated for activity modulation. Controlling the beating rate of cardiomyocytes is a pivotal technique in studies screening the pharmacological effects or toxicity of novel drugs in vitro.^[23]^ For instance, studying the beat rate-dependent actions of drugs requires cardiomyocyte activity modulation over a wide frequency spectrum.^[24]^ Moreover, in tissue engineering, electrical activity modulation during tissue development has been shown to influence differentiation and maturation in vitro.^[25]^

Typically, cardiac activity modulation is done electrically, using electrode arrays, or by optogenetic methods. While the latter technique requires genetic modification of the cells to introduce light-sensitive ion channels/pumps, the former is difficult to combine with currently emerging three-dimensional (3D) tissue constructs.^[26,27]^ Hence, mild thermal stimulation via GNPs^[28]^ provides an interesting alternative for both electrical or optogenetic stimulation modalities.

Thermal stimulation was shown to induce calcium oscillations in cardiomyocytes (leading to changes in contraction rate),^[28,29]^ remotely activate striated muscle cells via induced myotube contraction as well as modulation of related gene expression.^[30,31]^

In combination with 3D scaffolds, GNPs are excellently suited for optical, localized, temperature control. Few systems, however, have been developed to deliver localized thermal stimuli to cells or clusters of cells in 3D and integrated sensors for *in situ* remote read-out of the local temperature are lacking. In 3D hydrogels, thermal stimulation is often difficult to uncouple from mechanical stimulation due to hydrogel deformation.^[32]^ Previously we reported the integration of rhodamine-B infused silica particles as nanothermometers and gold nanorods (GNRs) into poly-ethylene glycol (PEG) hydrogels for cell actuation.^[32]^ The platform was a proof of concept as non-degradable PEG hydrogels are not well-suited for cell survival. Yet, to our knowledge, this is the only system available that features localized, remote optical temperature read-out in the 3D extracellular matrix. Various probes have been developed for nano-thermometry purposes,^[33–36]^ most often applied to intracellular temperature mapping^[37]^ or applied in cancer hyperthermia treatments,^[38,39]^ dynamic drug delivery,^[40]^ and wound healing.^[41,42]^ For in vitro thermal mapping, alternative techniques include near-infrared cameras^[43]^ or scanning thermal microscopy.^[44]^ Nevertheless, optical imaging using thermosensitive fluorescent NPs offers an excellent alternative for fast, real-time, high spatial resolution monitoring using imaging techniques.^[45]^ Optical temperature detection can be achieved through monitoring various temperature-sensitive luminescence properties such as intensity, lifetime, and spectral shifts, with high spatial and thermal resolution.^[36,46]^

In this work, the scaffolds are constructed by functionalizing paper-based microfibers with both GNRs and CdSe/CdS QDs and infused with collagen in order to support cell viability and function. In the following we demonstrate how light-based scaffold control can be achieved to deliver localized stimuli to modulate the activity of cultured cells, and combined with the optical imaging of culture/cell parameters to monitor tissue engineering constructs.

## Results and discussion

### 1. Paper-based microfiber scaffold: surface modification and biocompatibility

The fabrication steps of a synthetic tissue construct based on the smart nanocomposite scaffold are shown schematically in Figure 1A. First, Thorlabs lens cleaning tissue was modified to *in situ* introduce amine groups on the cellulose fibers using silane coupling chemistry according to the protocol described by Koga et al.^[47]^ Next, citrate-stabilized GNRs, prepared as described in Experimental Section, roughly 15 nm x 55 nm in size, were electrostatically adsorbed onto the positively charged fibers, followed by covalent attachment of the 9 nm CdSe/CdS, silica-encapsulated QDs. The QDs have a fluorescence emission peak at 640 nm and were water-solubilized through coating with an 8 nm silica shell, terminated with PEG ligands.^[48]^ This coating has been shown before to prevent cellular uptake.^[49,50]^ Finally, the paper tissue is coated with a mixture of 0.001% fibronectin, 2% gelatin, and 2% poly-L-lysine in water to improve cell compatibility. Figure 1 shows a brightfield microscopy image of the functionalized tissue in panel B and a fluorescence microscopy image in panel C.

**Figure 1.**
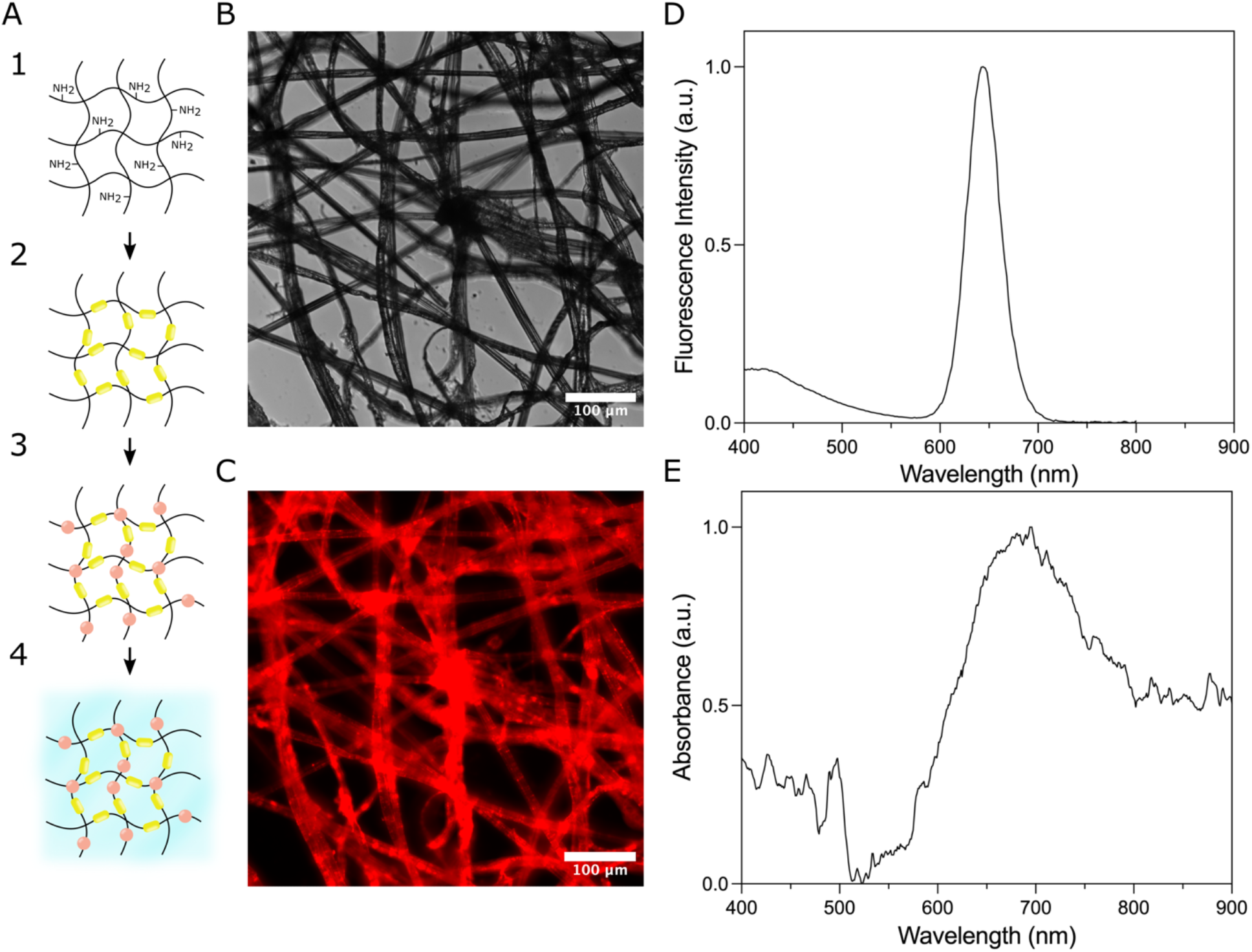
A) Schematic illustration of the steps in the fabrication of the paper nanocomposite scaffold: (1) chemical introduction of amine-groups, (2) electrostatic adsorption of GNRs (yellow), (3) covalent binding of QDs (red) and (4) collagen encapsulation. B) Brightfield microscopy image of cellulose paper at step 3. C) Fluorescence microscopy image of the QD-functionalized cellulose paper in panel B. D) Fluorescence emission spectrum of the QD functionalized the paper. E) Absorbance spectrum of the GNR functionalized paper.

The Thorlabs lens cleaning paper features a 3D network of fibers with an average diameter of (10 ± 2) μm and large pores, with an average diameter of (40 ± 25) μm, as determined using ImageJ’s particle analyzer from microscopy images as shown in Figure 1B. The highly porous structure mimics better the local cell microenvironment compared to 2D cell culture substrates and allows quick and efficient liquid absorption through capillary action.

We exploit the efficient wicking property of paper to firstly seed HL-1 cells onto the fiber mesh and, secondly, to facilitate the infusion of 20 μL of pre-gelled collagen solution. In this way, one can create cell-laden constructs with structural anisotropy, consisting of the functional, stiff cellulose fiber mesh surrounded by a secondary soft collagen matrix. Mechanical anisotropy is essential for the development and function of cardiac tissue.^[51,52]^ The collagen infusion also improves the scaffold attachment onto the glass coverslips for microscopy and, if desired, allows stacking multiple cell-laden cellulose sheets on top of each other, to create multi-layered structures.^[4]^ Collagen gelation effectively holds all layers in place. From here on we refer to this nanocomposite system as (QD-GNR-cellulose)/collagen.

The scaffold functions rely on the fact that the paper fibers can be easily functionalized to integrate stimulating and sensory NPs to remotely control and monitor specific cell characteristics with high spatial resolution. The optical properties of the particles used in this work, i.e., CdSe/CdS QDs and GNRs are shown in Figure S1 and S2 (Supporting Information), respectively. The functionalized paper also exhibits optical properties as shown in Figure 1D and 1E. GNRs have previously been incorporated into scaffolds for cardiac tissue engineering to modulate mechanical properties and/or increase the local electrical conductance,^[53]^ which are thought to increase cell alignment, elongation, and striation as well as induce higher expression levels of cardiac markers.^[54]^

In our scaffold, the GNRs serve as localized plasmonic heaters, with an LSPR peak at 750 nm. Under illumination by a 785 nm NIR laser, tuned to the absorbance peak of the GNR functionalized scaffold (see Figure 1E), local heat dissipation is induced locally in the scaffold through the excited LSPR waves. The QDs on the other hand, allow for simultaneous monitoring of the temperature through their temperature-dependent PL emission.

In the next sections, firstly, we discuss the organization of HL-1 cardiac muscle cells into the (QD-GNR-cellulose)/collagen nanocomposite scaffold, followed by the calibration of the thermal sensitivity of the scaffold and the QD detection of the NIR-induced local heating. Finally, in section 4, we demonstrate that the activity of cultured HL-1 cells can be modulated through NIR-induced plasmonic heating.

### 2. 3D cardiac tissue organization

HL-1 cardiac cells were seeded onto sterilized QD-GNR-cellulose scaffold of dimensions 0.5 cm x 0.5 cm and were allowed to sediment for 3 h at 37°C prior to the addition of extra growth medium. The next morning, cell-laden sheets were fixed within a SecureSeal 0.7 cm x 0.7 cm square chamber, attached onto a hydrophobic glass coverslip, and, subsequently infused with 40 μL of collagen gel. If desired, several cell-laden sheets could be stacked onto one another to create a layered structure.

After 5 days of culturing *in vitro*, the samples were fixated and stained with DAPI and phalloidin-AlexaFluor488 to image the cell cytoskeleton and nucleus, respectively (Figure 2). Furthermore, cell viability on 33 DIV cultured samples was investigated using the LIVE/DEAD viability assay under confocal microscopy (Figure S3, Supporting Information).

**Figure 2.**
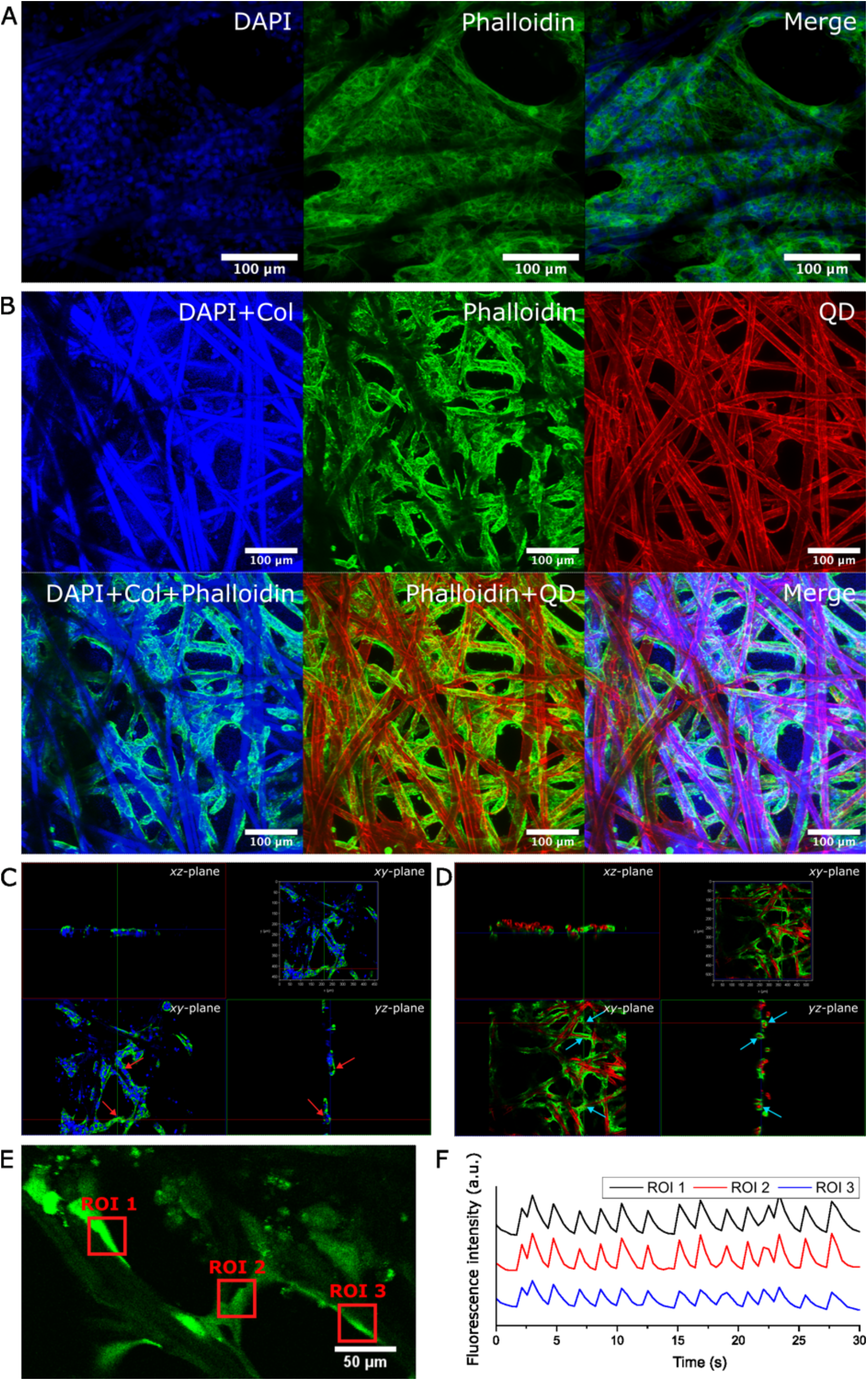
A) Confocal images with maximal projection 37.42 μm thick, DAPI/Phalloidin stained 5 DIV HL-1 culture inside a nanoparticle-free scaffold; B) Two photon confocal images with a maximal projection of 89 μm thick DAPI/Phalloidin stained 5 DIV old HL-1 culture inside a nanocomposite scaffold. Blue: DAPI staining and SHG signal of the cellulose and collagen fibers. Green: actin-fluorescent phalloidin staining. Red: QD fluorescence. C) 3D slice view of stained HL-1 culture inside a NP-free scaffold. Blue: DAPI staining and cellulose fibers. Green: Phalloidin staining. D) 3D slice view of HL-1 culture inside nanocomposite scaffold after 5 DIV culture. Green: Phalloidin staining. Red: QD fluorescence. E-F) HL-1 spontaneous activity calcium recordings in three ROIs displaying synchronized behavior, occurring after 2 days in culture. The cells were stained with Fluo-3 AM and a small area of the gel was imaged for 70 seconds, with an integration time of 433 ms per frame.

Figure 2 shows confocal fluorescence images of HL-1 cardiac cells cultured on both bare cellulose/collagen scaffold without NPs and QD-GNR-containing nanocomposite scaffolds, after 5 DIV (cell nuclei stained with DAPI can be visualized in blue, while the AlexaFluor488-phalloidin cytoskeleton is displayed in green). The cellulose fiber microstructure in the paper is also visible in the blue channel due to autofluorescence. Panels A illustrate the cell morphology in a 37.42 μm thick stack within a sample without NP-functionalization, while images in the Panel B are collected from an 89 μm thick stack in a QD-GNR-functionalized scaffold under two-photon confocal microscopy. The second-harmonic generation (SHG) signal (excitation at 900 nm) allows visualizing both the small collagen fibers and the much larger cellulose ones. QDs attached to the cellulose fibers are visible in the fluorescence channel in the panel B. On scaffolds without nanoparticle functionalization, the cells are homogeneously distributed on scaffold fibers and inside the interstitial collagen matrix, similar in morphology with the stem cell-derived cardiomyocyte cultures as reported.^[6]^ On the other hand, HL-1 cells cultured on QD-GNR-functionalized paper prefer to attach along the stiff cellulose fibers and elongate to form tubular constructs around them. The cross-sections shown in Panels C and D, also reveal the different cellular assembly, with more green actin staining localized along the fibers in the case of nanoparticle functionalized paper (see arrows). The observed architecture resembles more the one of the native myocardium, where layers of elongated cardiomyocytes combine into fiber-like structures to maximize force generation along the long axis of the tissue. *In vivo*, the extracellular matrix, (mechanically) supporting the tissue organization, contains collagen fibers aligned parallel to the cardiomyocytes to provide structural stability and tensile strength to which the cells adhere through a basement membrane featuring fibronectin and laminin.^[53]^

The HL-1 cells cultured in the nanocomposite scaffold exhibit spontaneous electrophysiological activity as evidenced by the recorded calcium transients shown in Panel F. Moreover, activity synchronization can be observed throughout the scaffold, illustrated in panel F by the calcium transients recorded in 3 distinct regions of interest (also see supplementary video 1). After prolonged culturing times (> 10 DIV), cells were still viable as confirmed by LIVE/DEAD staining, yet the continued cell proliferation caused cells to saturate the available space on the fibers, forcing expansion into the interstitial space filled by the soft collagen gel. Inside the collagen matrix, the cells assumed a more rounded morphology most likely due to the lower stiffness as compared to the nano-functionalized fibers. Stiffer filler materials, for instance, gelatin methacrylate with high crosslinking degrees, might be better suited for cardiac tissue formation.

### 3. Global temperature calibration of the nanocomposite scaffold

To read-out the local temperature changes, we calibrated first the temperature sensitivity of QD PL inside the nanocomposite scaffold. To this end, the nanocomposite scaffold was fixed on a glass coverslip and the temperature of the measuring chamber was varied cyclically between room temperature and 45°C.

Figure 3 shows the temperature-dependent fluorescence emission of a nanocomposite scaffold immersed in a 10 mM Hepes buffer at pH 7.4. In panel B, the buffer temperature is cycled between 21 °C and 45 °C, every 5 minutes, with the JPK Biocell™ temperature-control system, under constant illumination with 3.7 mW/mm^2^ green light (530nm). Over the course of a 40 min measurement, we observed (0.74 ± 0.002) % per minute photodarkening. The temperature-induced percentual change in PL intensity, displayed in panel C, is calculated from the fitted envelope functions, and remains constant throughout the measurement. The percentual fluorescence changes in response to temperature are calculated as

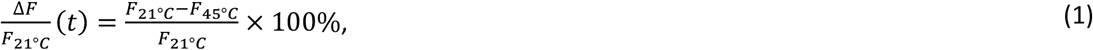

where *F*_21°*C*_ and *F*_45°*C*_ represent the fluorescence intensity values at 21 °C and 45 °C respectively, obtained from the envelope functions. A temperature increase of 24 °C induces a PL decrease of (40 ± 2) % (averaged over 3 different samples, with 3 measured areas per sample).

**Figure 3.**
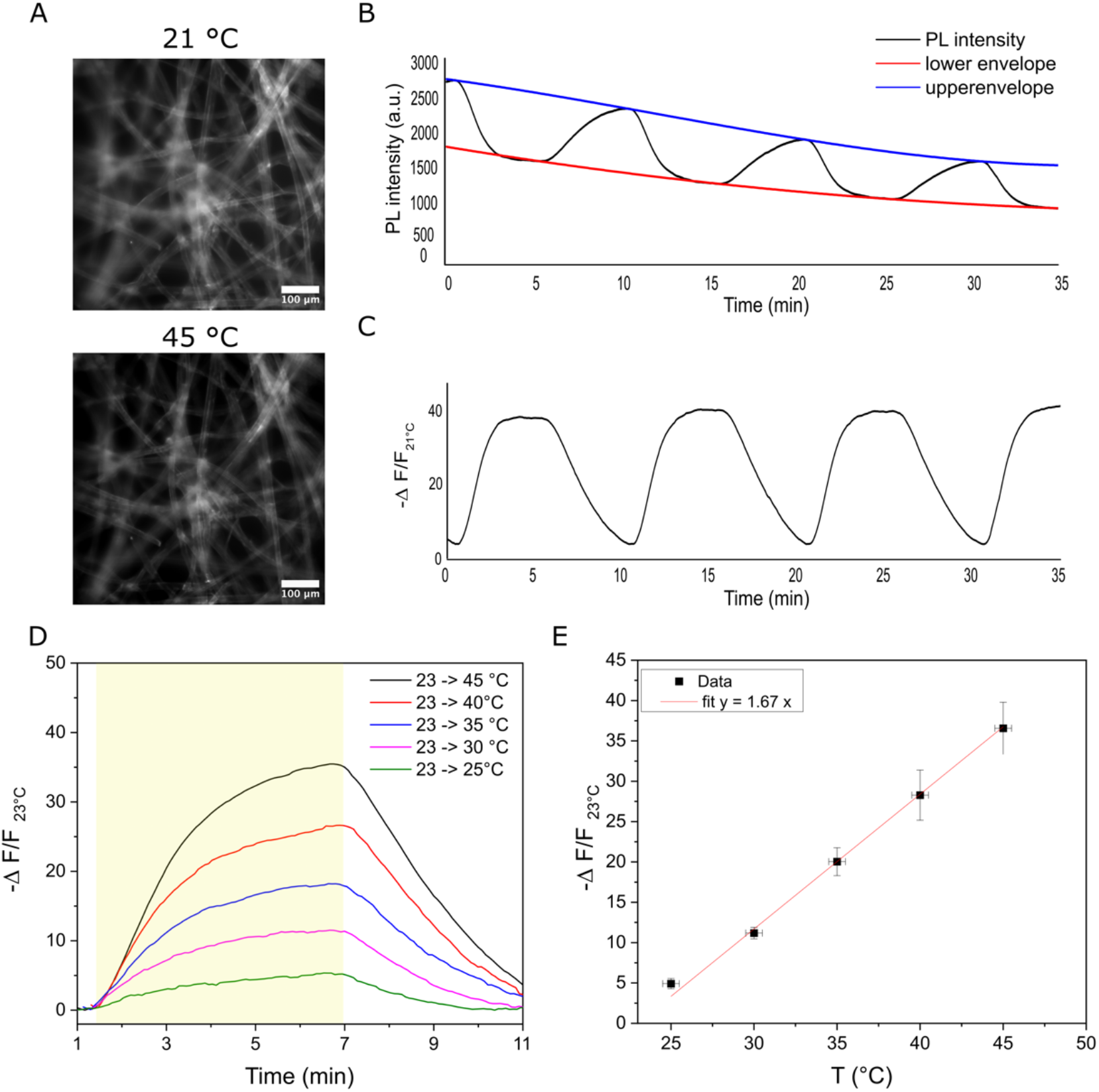
A) Fluorescence images of cellulose-GNR-QD fibers recorded at different bath temperatures (21°C and 45 °C, respectively). B) Temperature-dependent fluorescence changes in a nanocomposite scaffold upon cycling the bath temperature between 21 °C and 45 °C every 5 min and fitted envelope functions. The PL intensity is calculated as the integrated fluorescence intensity after background subtraction of each image (as in panel A) in a time series. C) Percentual change in QD PL as a function of time calculated using equation (1) for the data shown in panel B. D) Relative PL change in the nanocomposite scaffold for different temperature increments with respect to 23 °C as a function of time. The temperature is raised between minutes 1.5 and 7 and indicated in yellow. The percentual change is calculated similarly as was done in panels B-C, using the fitted envelope functions. E) Relative PL intensity change as a function of temperature, calculated from the data shown in panel D. The linear fit has a slope of (1.67 ± 0.09) %.

The time dependence of PL when increasing the bath temperature stepwise is shown in Figure 3D, for different temperature increments (see legend). The percentual changes in PL intensity 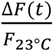 are plotted in function of the bath temperature in Figure 3E. In the range between 23 °C and 45 °C a linear trend is observed for temperature increments up to +22°C. From this data, we can extract the temperature sensitivity of the QDs in nanocomposite scaffold as being (1.67 ± 0.09) % per degree.

### 4. Plasmonic heat dissipation

Using a 785 nm NIR laser, focused on a specific region of the nanocomposite scaffold by a 10x objective, the local temperature can be raised through GNR plasmon heating. The focal spot is shown in Figure 4A and in this experiment has a diameter of 280 μm, corresponding to an illuminated area of roughly 61.5 · 10^3^ μm^2^. The local temperature can be controlled by tuning the NIR laser intensity (in our set-up between 0 and 90 mW). Figure 4b shows the changes in scaffold PL over time under intermittent NIR illumination (30 s period) with 10 mW (0.54 W/mm^2^), 50 mW (2.78 W/mm^2^), or 90 mW (5.11 W/mm^2^), respectively. In Figure 4b, the culture bath temperature is 23 °C and the applied NIR laser powers cause a local temperature increase of (1.2 ± 0.4) °C, (3.8 ± 1.5) °C, and (7.3 ± 1.9) °C for 10 mW, 50 mW, and 90 mW, respectively.

**Figure 4.**
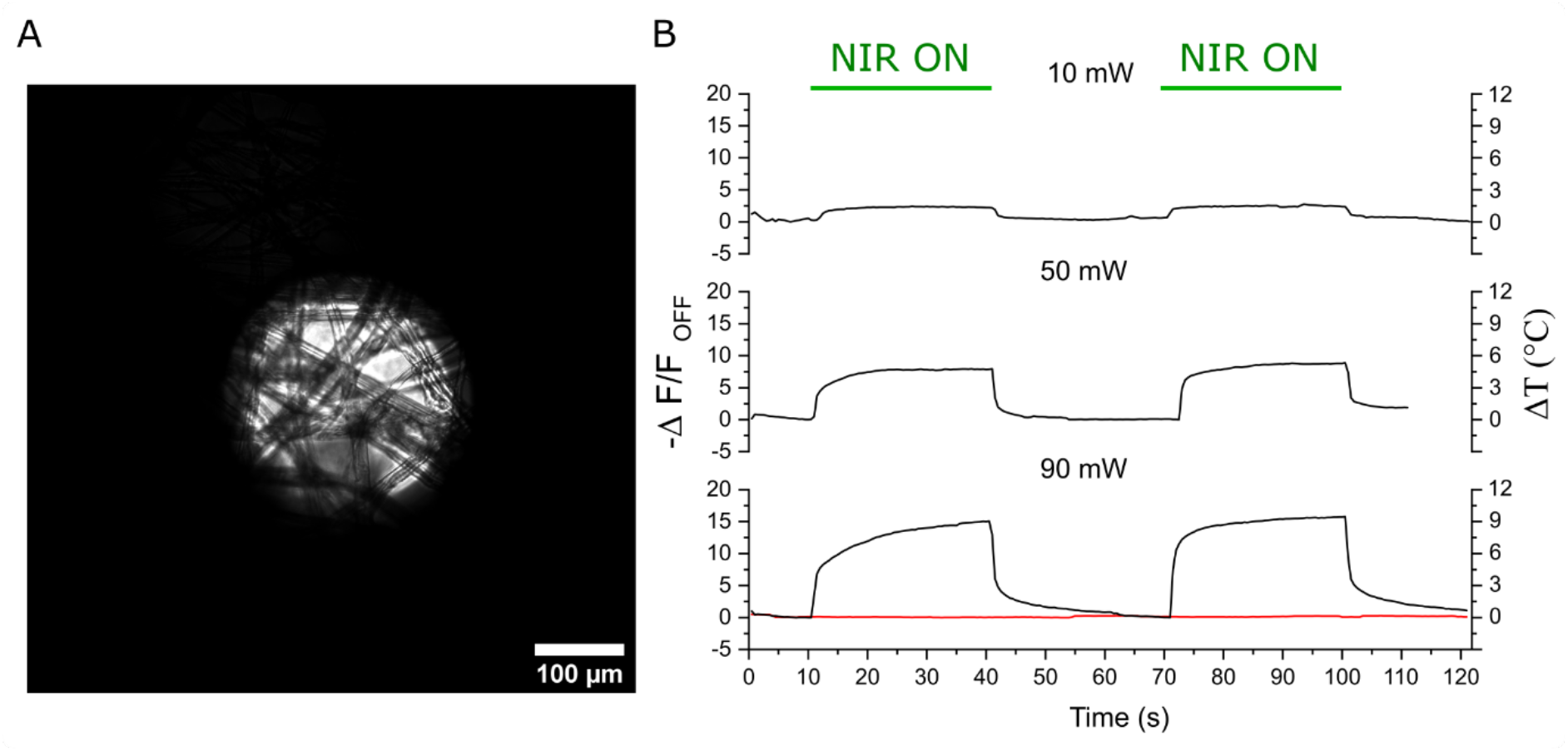
A) NIR illuminated area on a nanocomposite scaffold. B) Relative QD PL intensity changes for laser powers of 10 mW (irradiance 0.54 W/mm^2^), 50 mW (2.78 W/mm^2^), or 90 mW (5.11 W/mm^2^), respectively. The relative PL changes are calculated by 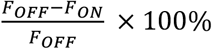 where F_ON_ and F_OFF_ are the averaged fluorescence intensities collected from a 100 μm × 100 μm square in the center of the NIR illuminated spot. The red trace shows the PL intensity when scaffolds without GNR functionalization are illuminated with 90 mW NIR.

### 5. NIR-controlled thermoplasmonic modulation of HL-1 activity

The effect of NIR-induced plasmonic modulation on the electrophysiological activity of HL-1 cardiac muscle cells has been studied on 3D cell cultures using the nanocomposite as the scaffold. For optical activity recordings, the cells are loaded with the fluorescent calcium indicator Fluo-3 AM to monitor the calcium transient as described in Experimental Section. Before performing plasmonic modulation on HL-1 cells, a reference experiment was conducted to check whether the NIR illumination affects the HL-1 electrophysiological activity in the absence of GNRs. As shown in Figure S3 (Supporting Information), the data shows no activity change upon NIR illumination without GNRs in the scaffold.

The GNR-mediated plasmonic modulation on HL-1 cell activity is shown in Figure 5. The modulation experiments were performed on cultures maintained at different bath temperatures, i.e., 25 °C, 30 °C and 37 °C (physiological temperature). In panel A, three representative calcium Fluo-3 fluorescence intensity traces are shown as a function of time. In each recording, the NIR laser is switched ON, to 90 mW (5.11 W/mm^2^) between 40 s to 80 s, as indicated. According to the calibration experiment discussed in the previous section, 90 mW NIR illumination causes a local temperature increment of (7.3 ± 1.9) °C, which in turn causes an increase in the activity rate of the cardiac cells. Panels A.1, A.2, and A.3 show examples of calcium activities in the HL-1 cells measured by the Fluo-3 calcium indicator intensity changes, upon NIR stimulation and for three base temperatures. For the traces shown, the temperature increase in trace A.1, from 25 °C to 32 °C, causes an increase in activity rate from 6.4 bpm to 16 bpm as evidenced by the Fourier transform, shown in the corresponding panel B.1. At a baseline of 30 °C, on the other hand, the rise in temperature causes an increased activity rate from 52 bpm to 64 bpm in trace A.2 (and B.2) while in trace A.3 (and B.3) at 37 °C baseline, plasmonic heating causes cell activity rate increase from 103.5 bpm to 105 bpm.

**Figure 5.**
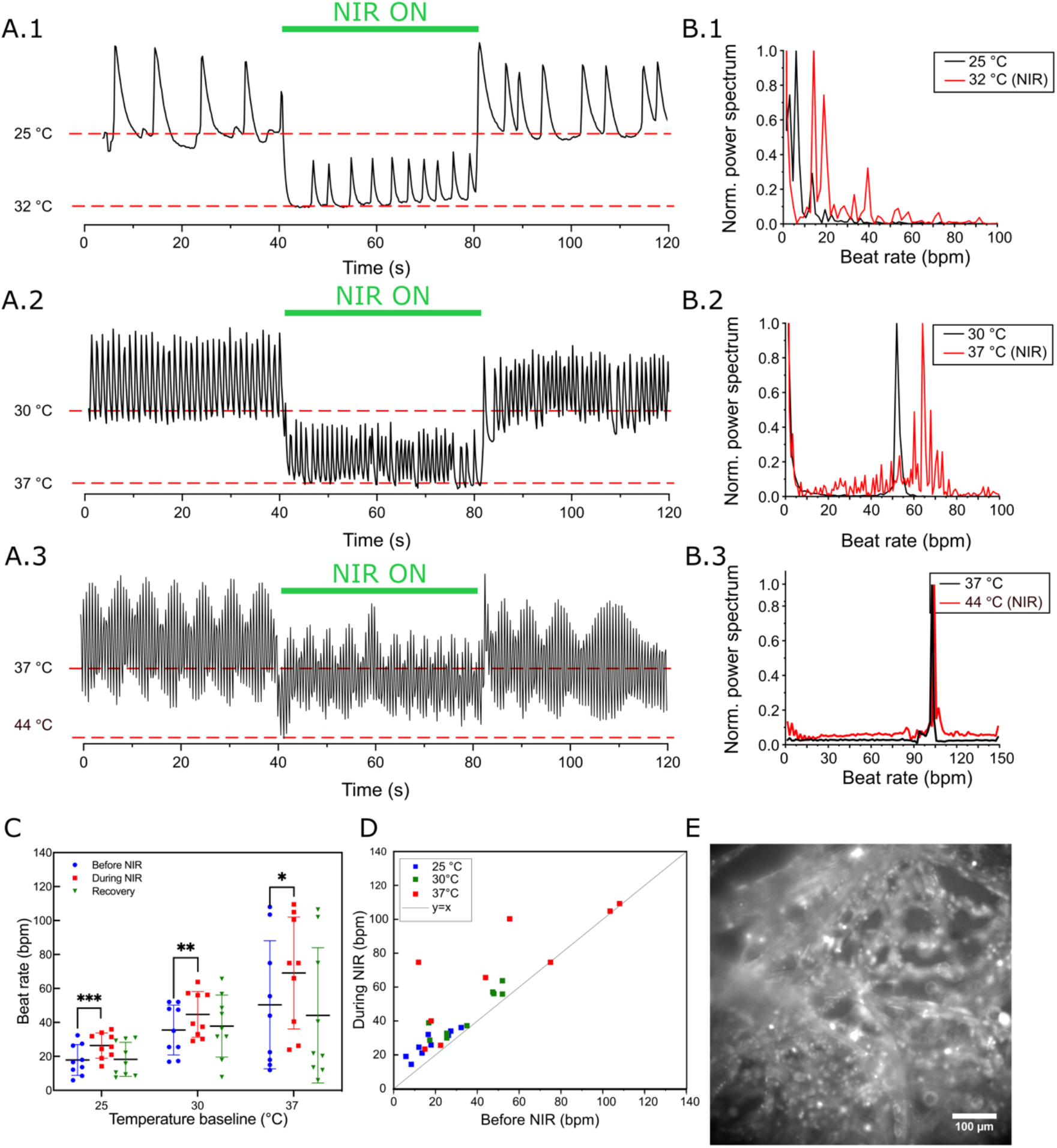
A) Fluo-3 calcium transients for 3 representative samples. Sample A.1 is maintained at a bath temperature of 25 °C, while samples A.2 and A.3 are at 30 °C and 37 °C, respectively. After 40 s the NIR laser is switched ON at 90 mW, causing a (7.3 ± 1.9) °C raise in temperature, locally. B) The corresponding Fast Fourier Transform (FFT) spectra of data in Panels A before (in black) and during plasmonic heating (in red). C) HL-1 beating rate (in bpm) before, during and after NIR (Recovery) plasmonic heating at different baseline temperatures. p-values were calculated from paired t-test, (*) p < 0.05, (**) p < 0.01, (***) p < 0.001, (****) p < 0.0001. D) Beating rates (in bpm) during NIR plasmonic heating in function of the initial activity rate (rate before NIR heating). E) Fluorescence microscopy image of Fluo-3 loaded HL-1 cells on the nanohybrid scaffold sample.

The statistical analysis of HL-1 activity rate variation in response to plasmonic modulation is shown in Figure 5C, comparing electrophysiological activity rate before, during and after NIR modulation (recovery), grouped in function of the base temperatures. In general, plasmonic heating consistently increases the beating rate of HL-1 cells for all base temperatures, which is consistent with our previous study on 2D cultured HL-1 cell samples.^[28]^ In Panel 5D, the cell activity rate is plotted during NIR modulation in function of the cell activity rate prior to NIR modulation. The data shows that at higher culture temperatures, cells have a higher beating rate and also a larger variability is observed in the beating rates across the culture. The data indicate that, at physiological temperature (37 °C), cells beating slower respond more to plasmonic stimulation compared to cells that beat initially at 80 bpm and higher.

The potential mechanism behind the temperature-dependent electrophysiological activity may be related to temperature-dependent ion channel activity.^[55,56]^ In addition to this, plasmon generation in GNRs produces not only a temperature increase but also strong electric fields and photocurrents, that all can stimulate different receptors in the cell membrane (e.g. through temperature-sensitive and voltage-gated ion channels) or by modifying the membrane capacitance, and as such contribute to the observed cell response. Future experiments should focus on varying the stimulation parameters in terms of NIR duration and intensity, eventually while blocking different types of membrane receptors, to understand the mechanisms of plasmonic modulation of cardiac activity.

### 6. Conclusions and outlook

In this work, we developed a functional, light-addressable, paper-based scaffold for cardiac cell culture that combines photothermal stimulation through plasmonic heating of GNRs with optical temperature and cell activity monitoring through the fluorescence of semiconductor QDs. The paper-based scaffold allows simple nanoparticle functionalization – not limited to the QDs and GNRs used in this study – and facilitates cell seeding due to effective fluid wicking. As such the nano-functional cellulose paper can be customized with a variety of nanoparticles and/or hydrogel systems. In our case, infusion of collagen, for instance, improves the scaffold attachment onto glass coverslips (for microscopy) and, if desired, enables stacking multiple cell-laden cellulose sheets on top of each other into modular design.^[57]^ One layer could then potentially accommodate cardiac cells while the next could host fibroblasts^[5]^ or enable the organization of endothelial cells for blood vessel organization.^[58]^ As such one can create thick, multicellular 3D synthetic tissues.

HL-1 cardiac muscle cells organized into oriented tubes around the stiff cellulose fibers in the paper and displayed synchronized electrophysiological activity as measured through calcium oscillations. Proof-of-concept experiments showed that plasmonic heating can accelerate the activity rate of the HL-1 cells depending on the initial temperature. The scaffold can be easily adapted depending on the cellular system under investigation, through the choice of paper to feature different fiber diameters or various degrees of fiber alignment for the creation of highly anisotropic cardiac patches. Moreover, due to the ease of functionalization, it is compatible with a wide range of nanomaterials that can act as thermal heaters or temperature-sensitive probes. And, additionally, using two-photon (confocal) microscopy can enhance the temperature sensitivity of QD ensembles, since the temperature induced changes in the two-photon PL are up to threefold larger than for single-photon fluorescence.^[59,60]^

Future experiments should study the exact experimental conditions that elicit the observed cell responses, i.e. increasing beat rates. In first instance, this can be done by tuning the stimulation parameters (NIR intensity and illumination duration, GNR concentration, etc.). Moreover, rather than increasing the beat rate with respect to the spontaneous activity rate, applications would benefit from actively pacing the beat rate. To this end, the use of short, (sub)millisecond NIR pulses at well-defined frequencies can be investigated for AP triggering.^[29]^

We believe that such platforms can provide powerful tools for developing in vitro model systems of cardiac arrythmia and screening the rate dependent effects of novel pharmacological products. Moreover, such a platform can be instrumental in studying the effect of temperature on cardiomyocyte activity.

## Experimental Section

### 1. Materials

All materials were used as received without any further purification. Cetyltrimethylammonium bromide (CTAB), Poly-L-lysine hydrobromide (PLL), 1-ethyl-3-(3-dimethylaminopropyl)-carbodiimide (EDC), sulfo-N-hydroxysulfosuccinimide (NHS), Pluronic F-127, dimethyl sulfoxide (DMSO), HL-1 cardiac muscle cell line, Fetal Bovine Serum (FBS), trypsin/EDTA, soybean trypsin inhibitor, gelatin, fibronectin, Triton X-100, norepinephrine, Grace Bio-labs SecureSeal adhesive sheet, trimethylchlorosilane, monomeric collagen solution from bovine skin, paraformaldehyde, DAPI dihydrochloride, calcein-AM and ethidium homodimer were purchased from Merck. Tri-sodium citrate was obtained from Acros Organics, (3-Aminopropyl)triethoxysilane (APTES) was obtained from ABCR, 4-(2-hydroxyethyl)piperazine-1-ethanesulfonic acid (HEPES), penicillin/streptomycin, Glutamax, Fluo-3 AM, Dulbecco’s Phosphate Buffered Saline (DPBS), phalloidin-AlexaFluor488 were purchased from Thermo Fisher Scientific. Ultrapure water (UPW, >18.2 MΩ) was used for all experiments and produced with a Sartorius Stedim Arium Pro VF system. Lens cleaning tissue (Thorlabs MC-5) was purchased from Thorlabs.

### 2. Nanoparticle synthesis

Citrate-stabilized gold nanorods (cit-GNRs) were synthesized following a seed-mediated protocol,^[61]^ and transferred from CTAB solution into sodium citrate solution following a capping agent exchange protocol.^[62]^ The detailed protocol and characterization have been reported in our previous work.^[28]^

Silica-encapsulated CdSe/CdS QDs (Ø 9 nm + 8.5 nm SiO_2_) with a PEG surface coating, were prepared by Aubert et al.^[48,49,60]^ Briefly, the QDs were first synthesized through the “flash” method followed with a carboxylated silica encapsulation procedure and PEG termination (9-12 PE units), characterization data can be found in Figure 1.

### 3. Nanohybrid scaffold preparation

Thorlabs lens cleaning tissue was cut into squares of 0.5x0.5cm and immersed in 10 mL of a 1% APTES solution in 80% (v/v) ethanol/water for 2h. The suspension was transferred to a single-neck, round-bottom flask, and the solvent evaporated using a rotary evaporator at 70 mbar, 40°C. The dried, silanized tissue fragments were heated to 110°C for 1h, rinsed 3 times with ethanol, and finally resuspended in distilled water for further functionalization.

The silanized tissue scaffolds were first functionalized with cit-GNRs through electrostatic adsorption. Briefly, APTES functionalized tissue scaffolds were immersed in 5.8 × 10^−10^ M cit-GNR solution overnight, followed by a rinsing with ultrapure water and drying with nitrogen gas.

Next, silica-encapsulated CdSe/CdS QDs were covalently bound to the APTES-modified GNR-cellulose scaffold via EDC/NHS chemistry. Briefly, 250 μl of 1 × 10^−7^ M QD suspension was mixed with 1 mL of 5 × 10^−2^ M MES buffer (pH 6.5) containing 2 × 10^−3^ M EDC and 2 × 10^−3^ M NHS. After 15 min, the cellulose-GNR sheets were immersed in the 2 × 10^−8^ M EDC-NHS activated QD solution for 4h and afterward rinsed twice with ultrapure water for 5 min. The functionalized QD-GNR-cellulose scaffolds were sterilized by immersion in 70% ethanol for 10 min followed by rinsing with DPBS, then they were coated with cell-adhesive motives to improve biocompatibility by sequential immersion in (1) a 0.02% (w/w) gelatin, 0.0005% fibronectin in water for 1h and (2) 1 mg/ml PLL in 100 mM borate buffer, pH 8.5. The final construct was rinsed twice with DPBS before use for cell culture.

### 4. Cell culture

HL-1 cells were purchased from Merck and cultured according to the standard cell culture protocol provided. Cells were maintained in Claycomb medium supplemented with 10 % FBS, 100 U/mL penicillin/streptomycin, 1 × 10^−4^ M ±-Norepinephrine and 2 × 10^−3^ M Glutamax in a Binder incubator at 37°C and 5% CO_2_. The T25 culture flasks were precoated using 1 mL of 0.02% gelatin 0.005 mG mL^-1^ and fibronectin in water for >1h at 37°C, followed by a rinse with DBPS.

Three times a week, upon confluency, the cells were passaged 1:3. Following a rinse with DPBS without Ca^2+^ and Mg^2+^, the cells were dissociated through the addition of 1 mL of 0.05 % trypsin/EDTA for 1 min at 37°C, which was then replaced by 2 mL of 0.05% trypsin/EDTA for another 2 min at 37 °C. Cell detachment was verified by optical microscopy and 2 mL of soybean trypsin inhibitor was added to quench the enzymatic reaction. Cells were then centrifuged and resuspended 1:3 in 5 mL of new growth medium in a freshly coated T25 flask.

HL-1 cells were seeded on the nanohybrid scaffold prepared in Section 3, 200 μL of 10^6^ cells mL^-1^ cell suspension were seeded on top of the nanohybrid scaffold for 3h at 37°C. Subsequently, growth medium was added, and cells were left to adhere on the scaffold overnight. Finally, the cell-laden constructs were infused with 20 μL of a 2 m mL^-1^ collagen gel and fixed into a 0.7x0.7 cm square chamber designed from Grace Bio-labs SecureSeal adhesive sheet, which was mounted onto a hydrophobic glass coverslip (25 mm diameter and 0.16 mm thickness). The glass coverslips were hydrophobized by first exposing them to UV/ozone for 10 min, followed by gas phase silanization in a closed container using 20 μL trimethylchlorosilane in a nitrogen atmosphere.Afterward the coverslips were baked at 110°C for 10 min and rinsed with isopropanol and water. The collagen gel was prepared by neutralizing a 6 mG mL^-1^ monomeric collagen solution from bovine skin. To this end, 666 μL of collagen stock solution was mixed with 100 μL of PBS 10X, 131.5 μL ultrapure water (>18.2 MΩ), 102.5 μL of 0.1M NaOH and 1 mL PBS 1X.

### 5. Fixation and staining

The HL-1 cell samples were fixated by aspiration of the growth medium and incubation in a fixation solution containing 4 % paraformaldehyde and 0.2% Triton X-100 for 30 min at room temperature. The fixated sample was then rinsed 3x with DPBS to remove excess paraformaldehyde and incubated in 0.1M glycine in DPBS for 30 min to block unreacted aldehydes, followed by two washing steps of 5 min with DPBS.

Live/dead cell staining was performed using calcein-AM and ethidium homodimer. Calcein-AM and ethidium homodimer were dissolved in DMSO at a concentration of 1 × 10^−3^ M. Then 2 μL of calcein-AM and 2 μL of ethidium homodimer were dissolved in 1 mL of DPBS to obtain a staining solution. After 30 min of incubation with the staining solution, the latter was replaced by DPBS, and the sample was directly imaged using confocal microscopy.

For DAPI/Phalloidin staining, Alexa 488-labeled phalloidin was dissolved in methanol to yield a 40X 6.6 μM stock solution whereas DAPI dihydrochloride (Sigma-Aldrich) was dissolved in UPW to yield a 100X 250 μG mL^-1^ stock solution. The staining solution consists of 1X phalloidin and 1X DAPI in PBS with 1% bovine serum albumin. The fixated sample was incubated for 2 hours with the staining solution at room temperature and then washed twice with PBS for 5 min and stored in PBS at 4°C protected from light until imaging.

### Calcium imaging

Calcium imaging was performed using the calcium indicator Fluo-3 AM. 50 μG Fluo-3 was dissolved in 14 μL DMSO containing 20 % (w/w) Pluronic F-127 to yield a 1000X stock solution. The 1X staining solution was then prepared by dilution with HEPES buffer (pH 7.4) and samples were incubated for 30 min with 500 μL staining solution, rinsed twice with DPBS and left for an additional 30 min in HEPES buffer (pH 7.4) before performing microscopy.

### 7. Fluorescence microscopy and confocal microscopy

For brightfield imaging and calcium imaging, an inverted fluorescence microscope (Olympus IX81) integrated with JPK Biocell™ temperature-control system was used. The calcium dye Fluo-3 emission was recorded using a U-MWIB3 Olympus filter cube, containing a 460-495 nm bandpass excitation filter, a 530 nm bandpass emission filter (Chroma, ET 535/30) and a 505 nm dichroic mirror. Excitation light was provided by an XCite white light source coupled to the backport of the microscope. The QD fluorescence was recorded using a U-MWIG3 Olympus filter cube, containing a 530 bandpass excitation filter, a 575 nm IF long pass emission filter, and a dichroic mirror with a cut-off of 570 nm. A 20x objective lens (Olympus LUCPLFLN20XRC, NA 0.45) was applied for fluorescence collection and a Hamamatsu ORCA-Flash 4.0 v2 scientific CMOS camera was used for image acquisition with an exposure time of 200 ms.

Confocal microscopy and multiphoton imaging were performed for Live/dead assay and DAPI/Phalloidin staining on Leica SP8 dive microscope and a custom-made microscopy setup consisting of an upright Olympus BX61 W1 microscope coupled to a tunable Insight DS femtosecond laser (a wavelength of 900 nm was used in this work). The fluorescence emission was collected using a 25x water immersion objective lens (HC FLUOTAR L 25x/0.95 W VISIR).

### 8. NIR-induced plasmonic heating

For NIR actuation of the nanohybrid tissue, the same set-up was employed as described in the work of Yu *et al*.^[32]^ Briefly, samples were also placed in a JPK Biocell™ temperature-controlled imaging chamber (Bruker, UK) which was mounted on the Olympus IX 81 inverted microscope. A 785 nm collimated NIR laser was fiber-coupled to a 10x objective (Olympus UPLANFL10X, NA 0.3) mounted in an ‘upright’ position above the sample stage.

## Acknowledgments

The authors thank prof. dr. Zeger Hens and dr. Tangi Aubert for kindly supplying the silica coated CdSe/CdS quantum dots. F. Guo acknowledges the financial support from the China Scholarship Council (CSC). S. Jooken acknowledges the financial support by the Flanders Research Foundation (FWO) 1SC3819N. C. Bartic, W. Thielemans, and O. Deschaume acknowledge the financial support through FWO grant G0947.17N and KU Leuven research grants C14/18/061 and C14/23/093.

## Author contributions

Conceptualization, C.B., F.G., S.J and O.D; methodology, F.G. (plasmonic NIR actuation), S.J. (nanocomposite scaffold and cell culture) and W.Y. (optical set-up); Validation: F.G., A.A.; Formal analysis, S.J., F.G and A.A.; Investigation, F.G. (NIR actuation experiments, cell culture, histological staining, confocal fluorescence microscopy and sample preparation for cell actuation), A.A. (cell culture, histological staining, confocal fluorescence microscopy and sample preparation for cell actuation) S.J. (fabrication of the nanocomposite scaffold); Data curation, F.G., S.J., O.D. and C.B.; Writing – original draft, S.J. and F.G.; Writing – review and editing, C.B., F.G, O.D., A.A. and W.Y., Visualization, S.J., A.A. and F.G.; Supervision; C.B.; Project administration, C.B.; Funding acquisition, C.B., W.T., O.D., F.G. and S.J. All authors have given approval for the final manuscript version.

## Supporting information

Fluorescence spectra and TEM of CdSe/CdS QDs, absorbance spectrum and AFM of GNRs, Live/Dead assay, blank NIR experiment, supplementary video 1 (HL-1 electrophysiological activity on nanocomposite scaffold after 3 DIV culture), supplementary video 2 (HL-1 thermoplasmonic modulation on nanocomposite scaffold after 3 DIV culture, 4X speed).

## Table of Contents

In this work, the design and fabrication of a light-addressable, paper-based, nanocomposite scaffold for optical modulation and read-out of in vitro grown cardiac tissue is reported. This scaffold consists of plasmonic nanoactuators and quantum dot nanothermometers enable modulating cardiomyocyte beating rate while monitoring the local temperature changes with single-cell resolution.

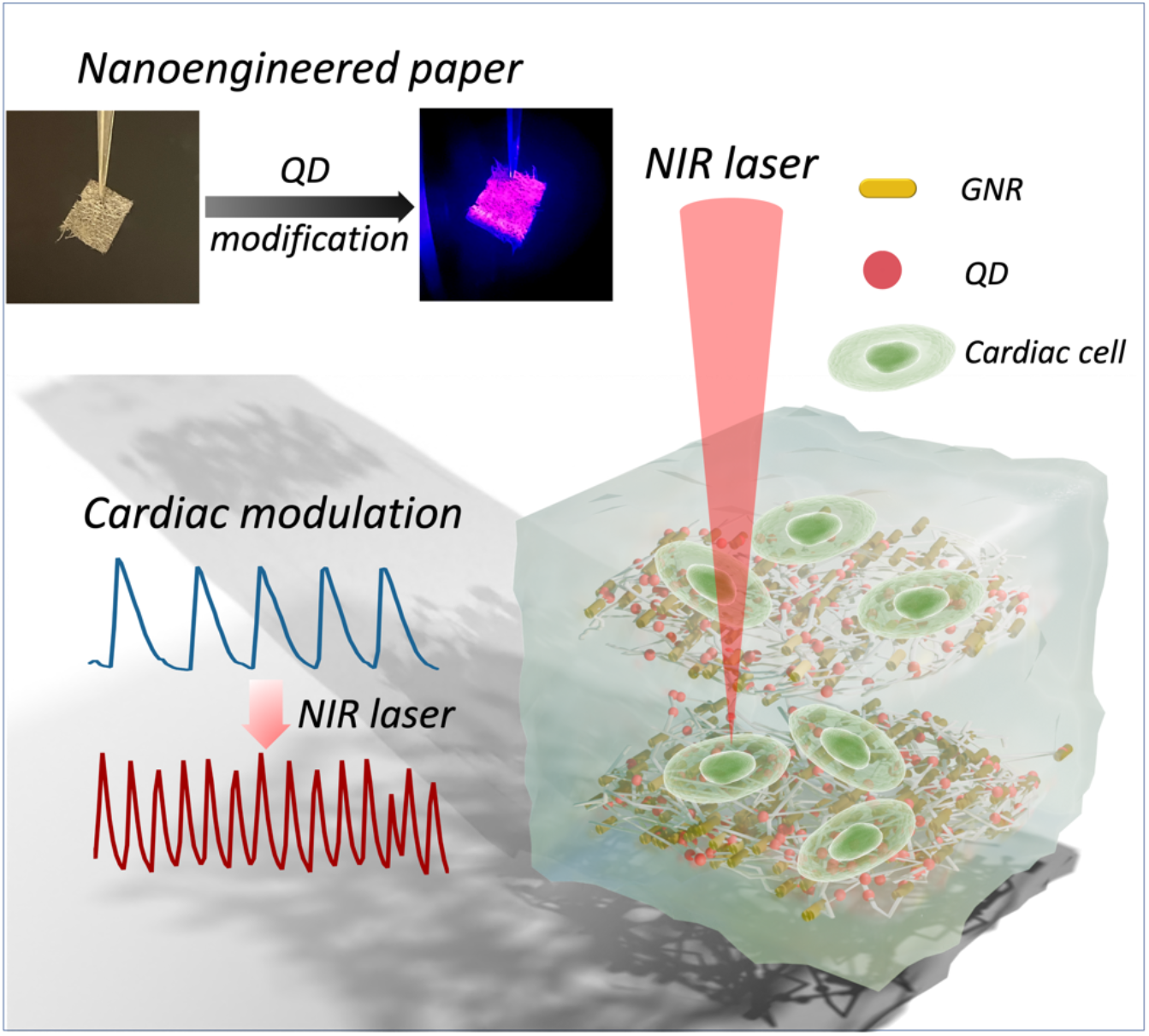

