## Supporting information for "Optically active, paper-based scaffolds for 3D cardiac tissue engineering"

**Overview of supporting information**

**Figure S1 Fluorescence emission spectrum and TEM characterization of the CdSe/CdS QDs**

**Figure S2 Absorbance spectrum and AFM characterization of the GNRs**

**Figure S3 Live/Dead assay on 33 DIV cultured HL-1 sample**

**Figure S4 Blank NIR experiment**

**Supplementary video 1 HL-1 electrophysiological activity on nanocomposite scaffold after 3 DIV culture**

**Supplementary video 2 HL-1 thermoplasmonic modulation on nanocomposite scaffold after 3DIV culture, 4X speed**

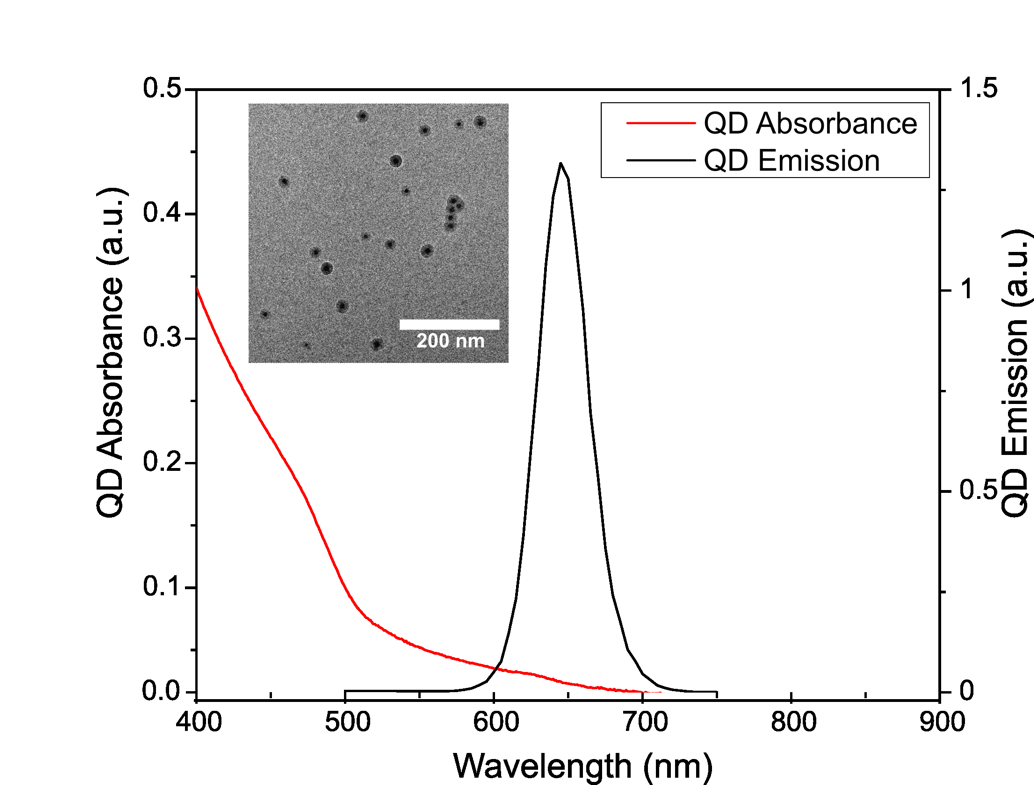

*Figure S1. Fluorescence emission spectrum and TEM characterization of the CdSe/CdS QDs.*

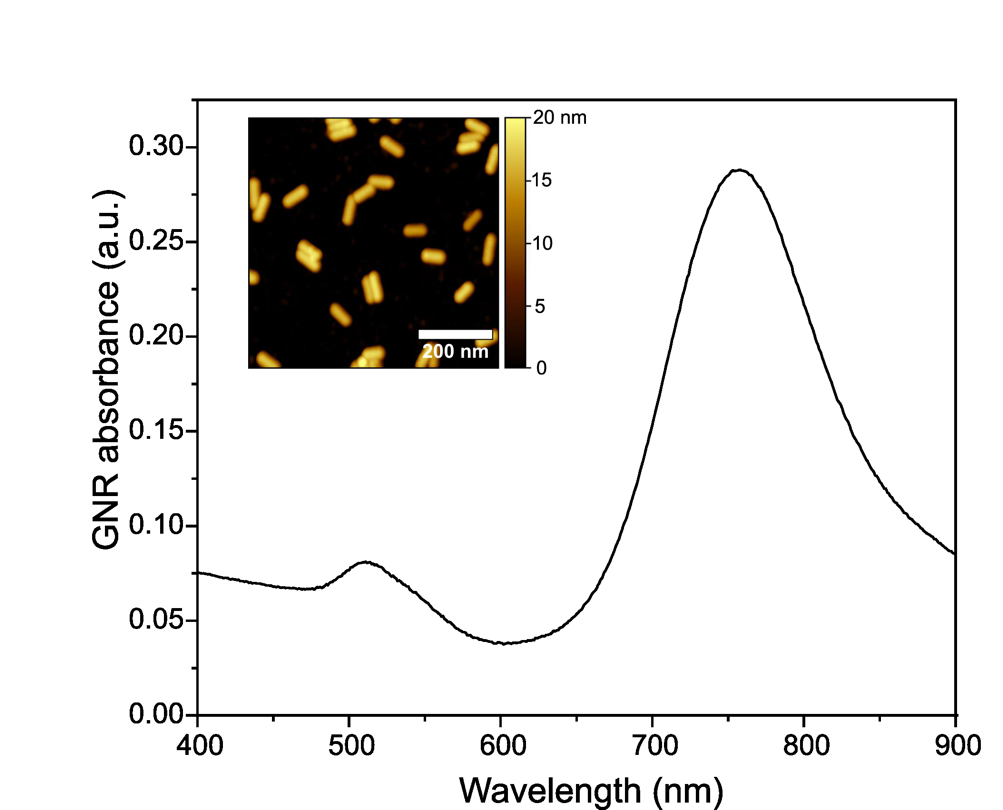

*Figure S2. E) Absorbance spectrum and AFM characterization of the GNRs.*

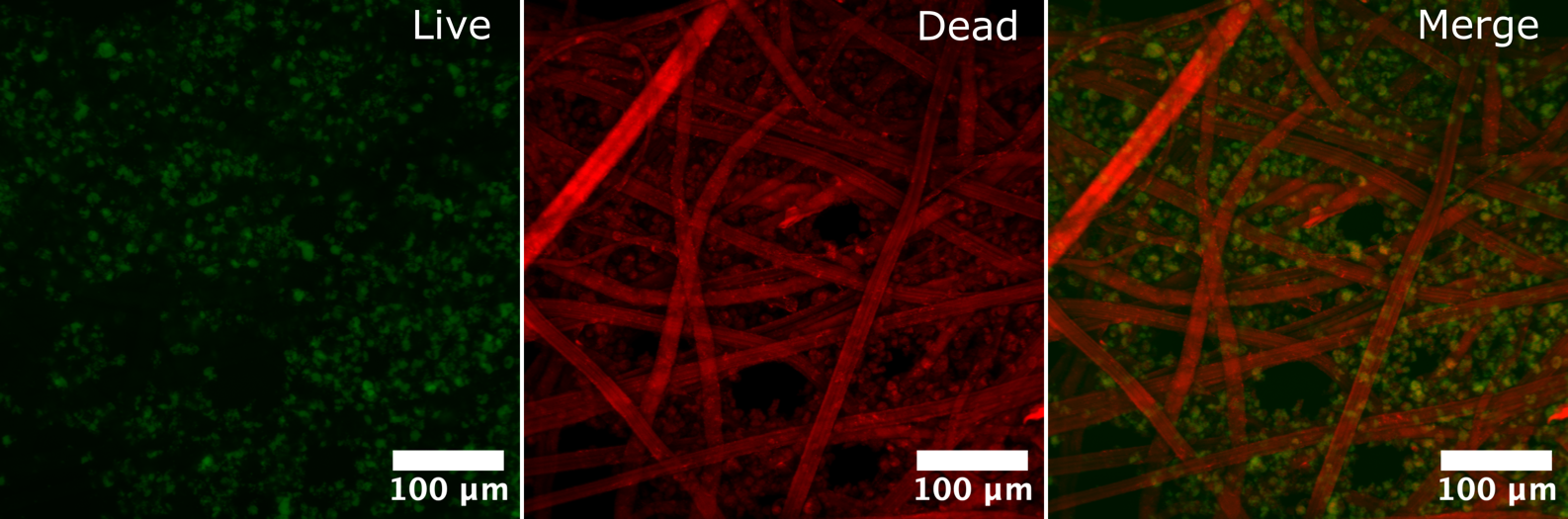

*Figure S3. Live/Dead assay on 33 DIV cultured HL-1 sample. Green: Calcein-AM stained live cells. Red: QD modified fibers and ethidium homodimer stained dead cells.*

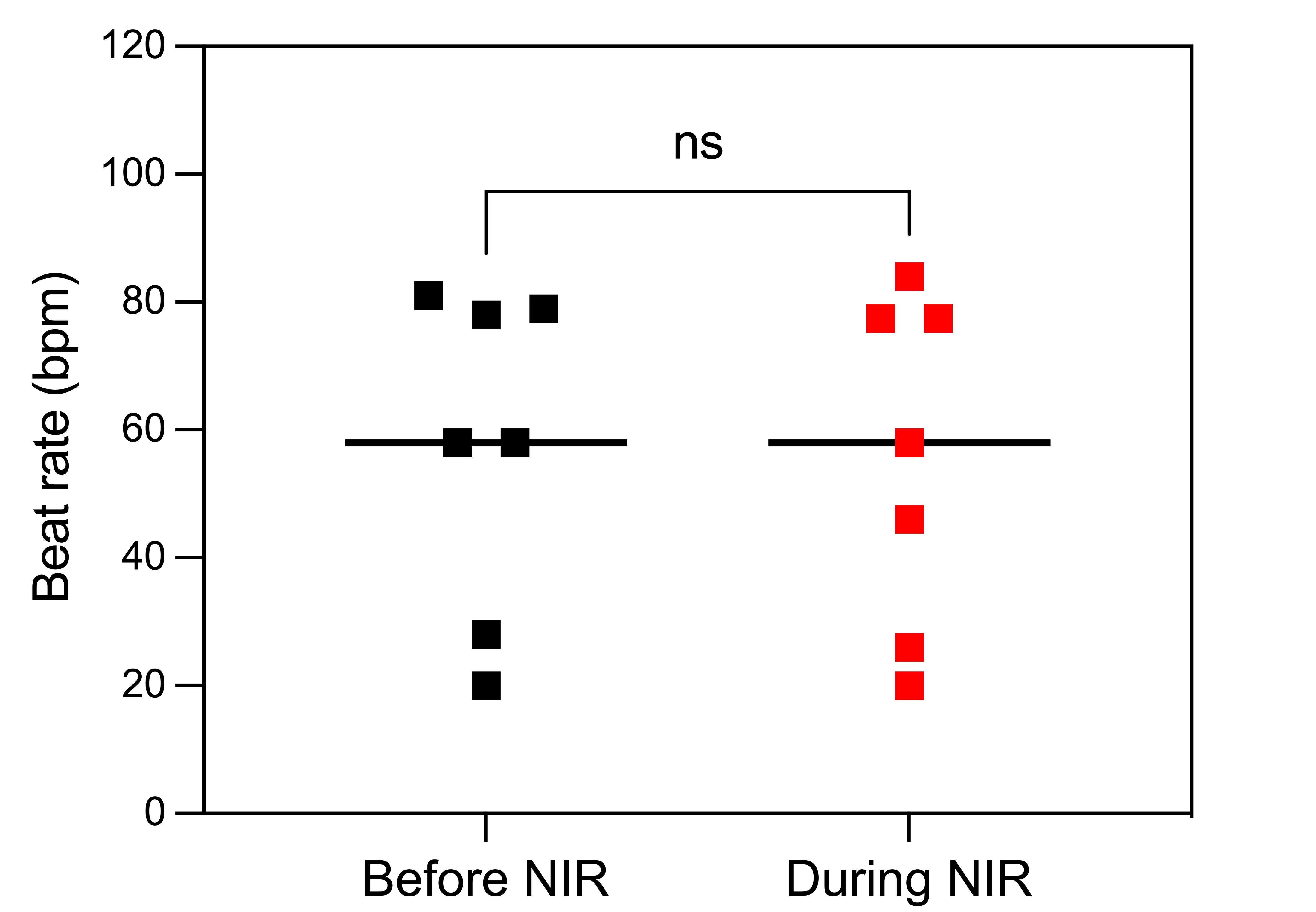

*Figure S4. Statistical results of 90 mW NIR stimulation on 5 DIV cultured HL-1 cells on cellulose scaffold without GNR functionalization, p-value calculated from paired t-test, (ns) not significant, N = 7.*
